# Exploring muscle recruitment by Bayesian methods during motion

**DOI:** 10.1101/2024.02.06.579136

**Authors:** M Amankwah, A Bersani, D Calvetti, G Davico, E Somersalo, M Viceconti

## Abstract

The human musculoskeletal system is characterized by redundancy in the sense that the number of muscles exceeds the number of degrees of freedom of the musculoskeletal system. In practice, this means that a given motor task can be performed by activating the muscles in infinitely many different ways. This redundancy is important for the functionality of the system under changing external or internal conditions, including different diseased states. A central problem in biomechanics is how, and based on which principles, the complex of central nervous system and musculoskeletal system selects the normal activation patterns, and how the patterns change under various abnormal conditions including neurodegenerative diseases and aging. This work lays the mathematical foundation for a formalism to address the question, based on Bayesian probabilistic modeling of the musculoskeletal system. Lagrangian dynamics is used to translate observations of the movement of a subject performing a task into a time series of equilibria which constitute the likelihood model. Different prior models corresponding to biologically motivated assumptions about the muscle dynamics and control are introduced. The posterior distributions of muscle activations are derived and explored by using Markov chain Monte Carlo (MCMC) sampling techniques. The different priors can be analyzed by comparing the model predictions with actual observations.

## 1 Introduction

Understanding the subtle interplay between the human central nervous system (CNS) and the musculoskeletal system (MS) to generate controlled movements to accomplish specific tasks is one of the central problems in neurophysiology as well as in biomechanics. The musculoskeletal system is characterized by muscle redundancy, whereby the number of muscles actuating a desired movement exceeds the number of degrees of freedom, implying that the desired movement can be accomplished in multiple alternative ways. While this redundancy is crucial for the system to work seamlessly under changing external conditions without being too sensitive to perturbations, it also raises the question of what strategy, if any, the human body follows to attain the desired goal. The reductionist approach assumes the existence of a cost function whose minimizer corresponds to the preferred muscle activation pattern, while the uncontrolled manifold hypothesis assumes that the activation pattern is an approximate optimizer among all possible activations respecting the physiological bounds. The uncontrolled manifold approach is more flexible and can account for patterns effectuating movements in which optimal activation is not possible, as may be the case for non-healthy, non-adult or aged subjects, or tasks performed in challenging environments, for example walking on slippery ground. Exploring the continuum of possible solutions to the muscle redundancy problem can be done by sampling techniques, including Bayesian Monte Carlo techniques [27, 28, 29]. For a comprehensive review of the different approaches that have been proposed in the literature, see [1].

As pointed out in the cited review article [1], a shortcoming of most solutions to the muscle redundancy problem is that the algorithms seek the solutions considering one time frame at a time. The reasons for this are twofold: From the purely computational point of view, solutions that are global in time require simultaneous treatment of the underlying underdetermined system over hundreds or more timeframes, increasing significantly the computational complexity. In addition, connecting the muscle activation patterns over timeframes requires a plausible model of how the central nervous system musculoskeletal (CNS-MS) complex selects the patterns over time under normal conditions. The model that we propose in this work assumes that under normal activation, i.e., when not performing maximal tasks, the body prefers patterns limiting the total yank [17], defined as the time derivative of the muscle forces. The present approach continues and extends the work in [27, 28] by putting the formalism in a Bayesian statistical context. More specifically, the intrinsic objectives and bound constraints are implemented as a priori information, while the information concerning equilibrium conditions obtained by motion tracking constitutes the likelihood: uncertainties in the tracking process as well as in translating the motion capture data into generalized joint force configurations contribute to the observation noise. In addition, we discuss previously proposed optimization strategies with a reinterpretation in the Bayesian framework providing a uniform framework for the comparison of different objective functions. The remainder of the article is organized as follows. In the next section we present a concise mathematical formulation of the muscle recruitment problem that is a basis for the likelihood model. Section 3 provides an outline of the Bayesian formulation. The full Bayesian model is developed in section 4, where we introduce prior densities encoding hypotheses about the physiological muscle recruitment principles. Section 5 presents the details of the computational engine, the MCMC sampling algorithm. The performance of the algorithm is demonstrated with computed examples in section 6.

## 2 Muscle activation problem

In this section, for completeness, we present a concise derivation of the forwards model relating the muscle forces and the movement that those forces generate, and state the related inverse problem whose solution is the main topic of this paper. The body is modeled as a link-segment system, and the muscle forces generate torques through a lever arm model at the joints with given degrees of freedom which translate into movements of the body. The forward problem consists of computing the trajectories of the joints from known muscle forces, while estimating the muscle forces from partial observations of the movement is the corresponding inverse problem of interest to us.

In biophysics and robotics, the inverse dynamic problem, or inverse structural dynamic problem seeks to find the generalized forces, or torques that applied to joints of a link-segment model of a human or an animal body or a robot effect a desired motion. For the sake of definiteness, we assume that the task of interest is level walking by a human, observed through a motion capture system that records the paths of fiducial markers attached on the surface of the body. Subsequently, the motion is encoded in terms of the degrees of freedom, typically a set of angular variables related to the joints of the link-segment model of the body. This translation is typically based on a weighted least squares process, fitting the dynamic model prediction based on the degrees of freedom to the observations, see, e.g., [8], and the manual of OpenSim for details of the fitting process. In the following, the dynamics of the link-segment model of the body is encoded in terms of the vector-valued function θ(t)*∈* ℝ^*m*^, 0 *≤*t *≤*T, where the m components of the vector represent the angular degrees of freedom associated to the joints characterizing the model of the body. It is assumed that the time course of the generalized coordinates, combined with geometric information including the inertia tensors of the limb segments and the reaction forces completely characterize the dynamics of the system.

To solve the inverse dynamic problem of finding the generalized forces, consider the Lagrangian function of the joint segment system, given by

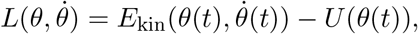

where E_kin_ is the kinetic energy of the system, and U is the potential energy [14]. The Lagrange equations

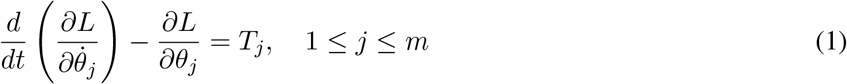

relate the generalized forces T_*j*_ and the equations of motion of the system. Assuming that the generalized coordinates are the joint rotation angles, the generalized forces represent the corresponding torques. By writing the kinetic energy as a quadratic form,

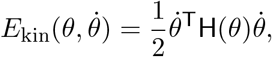

where H(θ) *∈* ℝ^*m×m*^ is the positive semidefinite (multi-body) inertia matrix of the system, it follows from (1) that the equations of motion can be expressed as

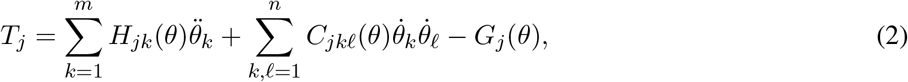

where C is the Christoffel symbol,

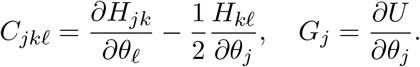

Here, the second term on the right accounts for centrifugal and Choriolis forces at the joints. These are the joint torques that the muscles need to effectuate to produce the observed trajectory. Since the time evolution of the generalized coordinates is approximately known, equation (2) allows the computation of the torques at times t_*ℓ*_, 0 *≤*ℓ *≤*N, denoted here by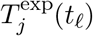. To express the torques in terms of forces, denote by 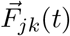 the vectors of forces of the muscles exerting a torque at the joint related to the jth generalized coordinate θ_*j*_, and let 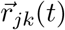 be the corresponding lever arm vectors. If 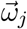 is the direction of the jth torque vector, we must have

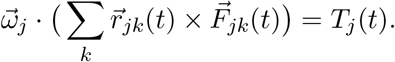

By collecting the scalar amplitudes of the forces of all n muscles involved in the system into a vector F(t) *∈* ℝ^*n*^, the torques in a vector T (t) *∈* ℝ^*m*^, and organizing the coefficients of the above equation into a lever arm matrix A(t) *∈* ℝ^*m×n*^, we arrive at the matrix equation,

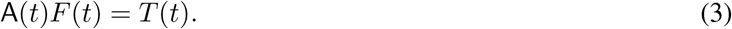

Denote the maximal (tetanic) forces of the muscles by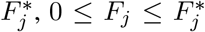, and define the muscle activations q_*j*_ by the formula

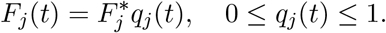

We express the equilibrium conditions (3) at the discrete times t = t_*ℓ*_ in matrix form as

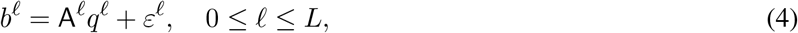

where b^*ℓ*^ *∈* ℝ^*m*^ represents the vector of estimated torques T ^exp^(t_*ℓ*_), q^*ℓ*^ = q(t_*ℓ*_) *∈* ℝ^*n*^ is the vector of activations at t = t_*ℓ*_,

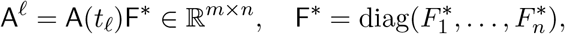

and ε^*ℓ*^ represents the uncertainties due to the estimation process of the inverse kinematic problem.

Equation (4) is the forward model describing how the muscle forces yield the generalized forces acting on joints to effect the observed motion. The corresponding inverse problem, which is the focus of this article, can be stated as follows:

### Muscle recruitment problem

Given the lever arm matrices A^*ℓ*^ at discrete time instances and the torques b^*ℓ*^ estimated with a given accuracy from the motion information, characterize the ensemble of possible muscle activations q^*ℓ*^ *∈* ℝ^*n*^, 0 *≤* ℓ *≤* L that are able to effectuate the observed movement, subject to additional suitable constraints.

In the following section, we discuss possible constraining conditions and develop a systematic Bayesian methodology to solve the muscle recruitment inverse problem.

## 3 Bayesian inverse problems: A recap

Before discussing the suitability and implications of different types of constraints, we provide a brief overview of the Bayesian methodology for inverse problems. For further details, we refer to monographs [4, 22].

Consider the problem of estimating an unknown quantity x *∈* ℝ^*n*^ based on the observation of a related quantity b *∈* ℝ^*m*^ as well as on a priori information about x. For the sake of clarity, we assume that the unknown and the observation are related to each other through a model

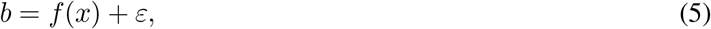

where f : ℝ^*n*^ *→* ℝ^*m*^ is known, and the vector ε *∈* ℝ^*m*^ accounts for both observation noise and uncertainties in the model. In the Bayesian framework, all unknown quantities are modeled as random variables regardless of whether the unknown represents a fully deterministic quantity or not: in other words, randomness in the Bayesian framework is an expression of lack of certainty about the value, regardless of the nature of the uncertainty. With that in mind, we interpret the equation (5) as a model relating the three random variables characterized by the respective probability distributions. We assume here that the variables x and ε are mutually independent. Assuming that the probability distribution of ε is defined through a probability density function π_*ε*_, that is

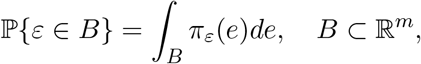

the probability density of **b** conditional on a known value of x is

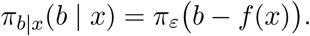

This density, referred to as the *likelihood density*, expresses the distribution of the observed values of b assuming that x is known and fixed. On the other hand, before observing b, we may have some information about the distribution of the unknown x which we encode in the *prior density* π_*x*_,

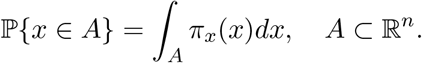

By the law of total probability, the *joint probability density* of the pair (x, b) is then given by the expression

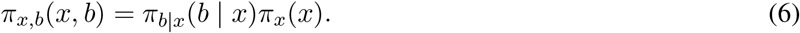

Reversing the roles of the variables x and b, the joint probability density of (x, b) can be expressed also as

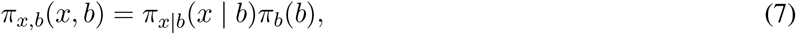

where π_*b*_ represents the marginal density of the variable b,

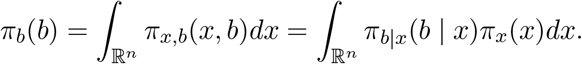

Equating the two equivalent representations (6) and (7) of the joint density naturally yields *Bayes’ formula*, relating the *posterior density* π_*x*|*b*_(x | b) to the likelihood and prior densities,

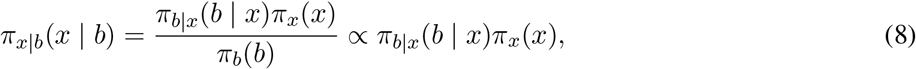

assuming that π_*b*_(b)/= 0. In the expression for the posterior, π_*b*_(b) represents a scaling factor that is unimportant in most of the analysis and is often omitted, hence the symbol “*∝*” indicating “proportional up to a scaling factor”.

Given the posterior density, it is common to summarize it with either the Maximum A Posteriori (MAP) estimate, or the Posterior Mean (PM) estimates,

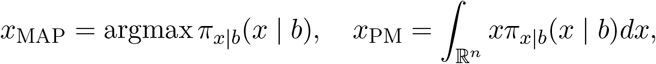

assuming that the above quantities are well defined. The estimate x_MAP_ is the solution of an optimization problem, while the computation of x_PM_ requires integration in ℝ^*n*^. In high dimensional problems, it is not feasible to integrate using numerical quadratures, and one has to resort to Monte Carlo techniques. A common approach is to use Markov chain Monte Carlo (MCMC) techniques to generate a large sample of realizations,

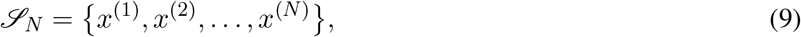

drawn from the distribution π_*x*|*b*_(x | b), then using the sample to estimate the posterior mean as

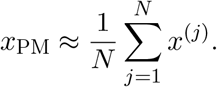

Observe that the sample *ℒ*_*N*_ provides much more information about the posterior distribution than just the posterior mean: For instance, we may approximate the posterior covariance matrix by

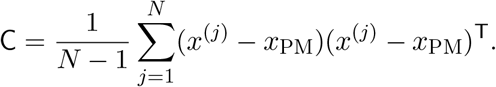

Moreover, the posterior (marginal) belief intervals of p% of the components x_*k*_, denoted by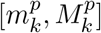, can be determined by sorting the components 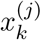 in increasing order,

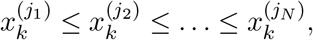

and discarding 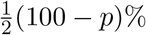 of the values from the top and from the bottom. More precisely, letting

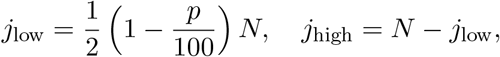

we define

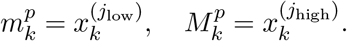

We end this section by outlining the MCMC sampling algorithm that will be used in the numerical calculations. Consider the current sample point x^(*j*)^. The Gibbs sampler algorithm provides a method to generate the next sample point x^(*j*+1)^ in the chain, drawn from the posterior probability distribution in the following way. For notational simplicity, let us denote the posterior density by π(x) = π_*x*|*b*_(x b), and let π(x_*k*_| x_1_, x_2_, …, x_*k−*1_, x_*k*+1_, …, x_*n*_) denote the one-dimensional posterior density of the component x_*k*_ obtained by keeping all components in the density π(x) fixed except for the kth one. The updating step of the Gibbs sampler consists of n random draws from onedimensional densities: Given the current sample point x^(*j*)^,

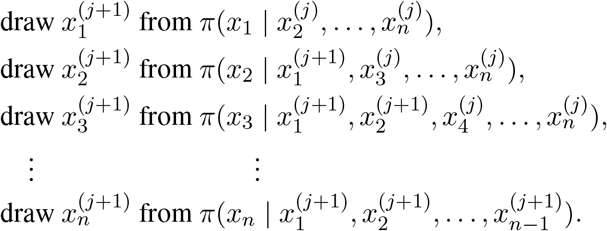

The implementation of the Gibbs sampler is, at least in principle, a straightforward task, requiring only random draws from one-dimensional densities. We will discuss the details that need to be addressed for a successful implementation in the context of computed examples.

## 4 Bayesian models for the muscle dynamics

In this section, we develop a family of Bayesian prior models designed to encode physiologically meaningful information about the muscle dynamics.

### 4.1 Likelihood model

In the Bayesian framework, we treat the equilibrium conditions (4) as observation models. For reasons of computational simplicity, we assume that the noise vectors ε^*ℓ*^ are mutually independent, and moreover, we model them as identically distributed scaled multivariate Gaussian white noise vectors,

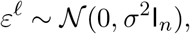

where σ > 0 is the noise level, and I_*n*_ is the n × n identity matrix. This yields the Gaussian likelihood model,

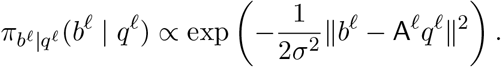

We organize the muscle activations into a matrix with columns indexed by the time instances and the rows by the muscles,

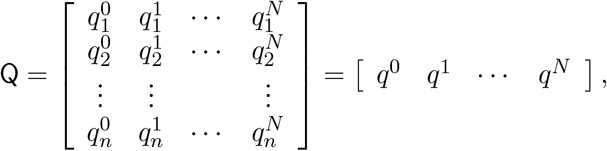

and we arrange the observations in a similar fashion into a matrix B *∈* ℒ ^*m×*(*N*+1)^. To vectorize the calculations, we stack the solumns of Q and B into the vectors

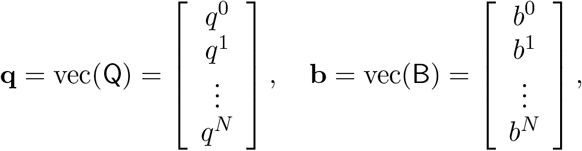

and we assemble the block diagonal matrix *𝒜*,

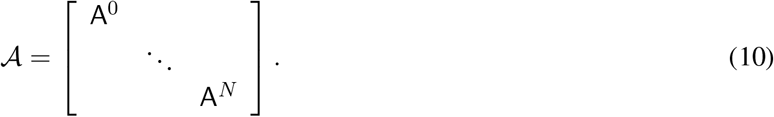

Assuming mutual independence of the identically distributed noise vectors ε^*ℓ*^, the likelihood of the time series of the generalized forces is of the form

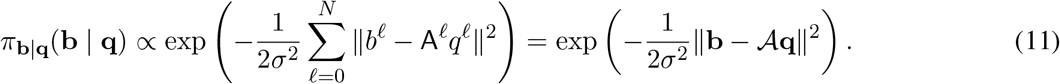

This likelihood model can be augmented by complementary information, for example electromyographic (EMG) measurements as indicated later in the discussion.

### 4.2 Prior models

In Bayesian analysis of inverse problems, the prior plays a crucial role, as it is a way to augment the observations accounted for in the likelihood with additional information that may be available. In underdetermined ill-posed inverse problems, where the data alone do not suffice to solve the problem in an unambiguous way, the prior can be thought of as adding constraints similar to regularization in the classical deterministic inverse problems framework [3]. In the following subsections we present several prior models, based on physiologically justifiable assumptions, that help explore the uncontrolled manifold in a more insightful manner.

#### 4.2.1 Box prior

Muscle forces must be non-negative and cannot exceed their presumably known tetanic values. These conditions can be expressed in terms of the muscle activations as bound constraints

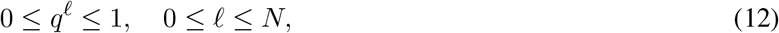

with the inequalities to be understood in component-wise sense. We encode these bounds into a *box prior* density of the form

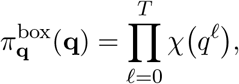

where χ is the multivariable indicator function of the unit hypercube in R^*n*^.

#### 4.2.2 Minimum activation prior

The information encoded in the box prior, while reducing considerably the range of possible activations, still allows a lot of redundancy in the determination of the force configurations, some of which may be physiologically justified under certain conditions, for example co-contraction forces at the joints. While simultaneous activation of agonist and antagonist muscles may occur, e.g., when joint stabilization is necessary, it is unlikely to occur under optimal conditions because it is energetically wasteful. It may be argued that the body may prefer activation configurations that minimize the need of energy metabolism, a hypothesis corroborated by EMG measurements [33, 6]. Following this principle, it is reasonable to hypothesize that the body seeks to minimize muscle activation. This is encoded in the *minimum activation* prior,

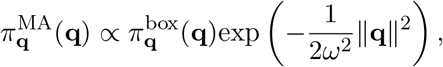

where the prior variance ω^2^ > 0 expresses how strongly the minimum activation principle is believed to be followed by the muscles. If there are reasons to believe that the task being performed requires co-contractions (e.g., level walking on ice), a larger variance value may be used to weaken the trust in the minimum energy principle.

#### 4.2.3 Minimum yank prior

The two priors introduced so far assume that the muscle activation patterns at the different times instances t_*ℓ*_ are mutually independent. This assumption may not be physiologically justified, because there are limitations for the speed at which an activation configuration can change. Assuming that the velocity at which the activation configuration takes place is a limiting factor, it is reasonable to hypothesize that in a normal continued activity such as level walking, the muscle forces do not change too abruptly in time. While in physics the time derivative of a force has no particular significance, in [17] the authors introduce the term ‘yank’ to refer to the time derivative of a muscle force. In physiology, yank is an important and useful concept: maximizing the yank improves a predator’s chances of reaching a target area, and the prey’s chances to escape. In [20] the authors discuss the problem of finding the cost function to be minimized to reduce the redundancies in muscle activations during healthy gait, highlighting the importance of the yank in sensorimotor systems, Furthermore, in support of their position they argue that mitigating the impact of external forces, identified with yank, could be detected by the cutaneous mechanoreceptors in the foot [24, 26], pointing towards the importance of the yank in the feedback mechanism in normal human walking. It has been suggested that controlling the yank is a way for the body to avoid injuries in leg tissues [30, 31], and in [21] it was shown that yank control may be motivated by energetic considerations because fast muscle cells use significantly more ATP than slow muscle cells.

Motivated by these considerations, we introduce a prior model favoring solutions with low yank between time frames over those with large yank. The yank of the kth muscle, defined as the time derivative of the force,

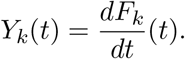

The average yank over the time interval [t_*ℓ−*1_, t_*ℓ*_] is given by

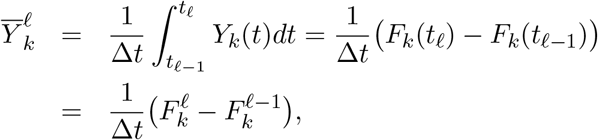

where F ^*ℓ*^ = F(t_*ℓ*_) *∈* ℝ ^*n*^. To formulate the belief about the yank in terms of a probability density function, for each muscle we write

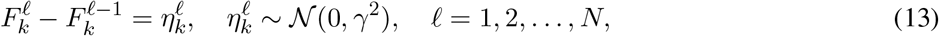

where we assume that γ *→*0 as Δt *→*0, guaranteeing continuity of the muscle forces. We can express condition (13) in vectorial form simultaneously for all muscles in terms of the activation vectors q^*ℓ*^ with the help of the diagonal matrix F^*\**^ of the tetanic forces, so that F ^*ℓ*^ = F^*\**^q^*ℓ*^. We write the component-wise equations (13) for each ℓ as

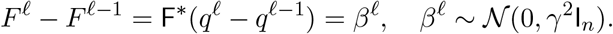

Next we consider all time instances together and write the matrix equation

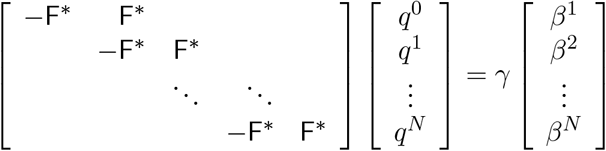

which can be expressed as

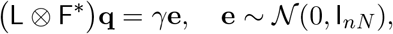

Where

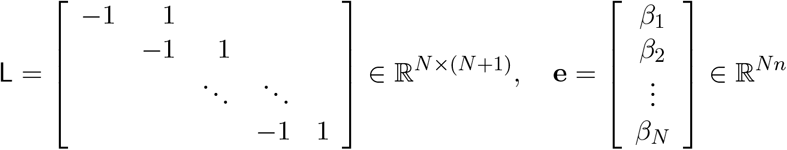

and *⊗* is the Kronecker product.

We are now ready to write the corresponding prior model for the paths **Q** as

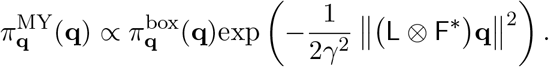

Observe that unlike the priors defined previously, the yank prior ties the time slice activations together to form paths, thus the realizations at different time slices are no longer mutually independent.

#### 4.2.4 Mixed prior

We end this section with a mixed prior model that interpolates the minimum activation prior and the yank prior. We define the prior density as geometric interpolation of the form

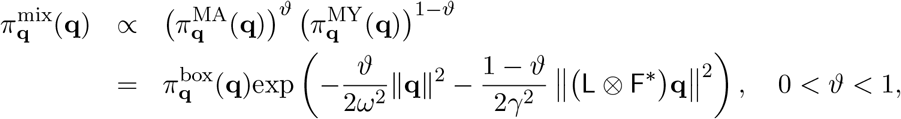

where the parameter ϑ determines the relative weight of the two priors. We remark that in the limit ϑ *→*0+ we have the minimum yank prior, while letting ϑ *→*1*−* we obtain the minimum activation prior. More generally, the priors can be combined as

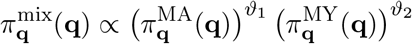

with mutually independent exponents ϑ_1_ and ϑ_2_. In our numerical simulations, we restrict the model to the former one to reduce the number of parameters.

## 5 Bayesian exploration of the posteriors

In this section we describe different ways to explore the posterior densities corresponding to the priors introduced in section 4. In the computed examples, since the number n of muscles exceeds the number m of degrees of freedom, the matrices A^*ℓ*^ *∈* R^*m×n*^ have a non-trivial null space, hence the posterior density for the box prior does not have a unique maximizer. Therefore, in the following discussion, MAP estimates are considered only for the minimum activation, minimum yank, and mixed priors.

### 5.1 MAP estimates by optimization

One popular way to summarize the posterior density is by means of the realization with “highest probability”, i.e., the MAP estimate. We start by considering the posterior distribution corresponding to the minimum activation (MA) prior. Since in this case the time slices are mutually independent, it suffices to consider single time slices. In the following, to simplify the notation we omit the superscript referring to the time instance, and write A = A^*ℓ*^, b = b^*ℓ*^ and q = q^*ℓ*^ for t = t_*ℓ*_.

With these notations, the posterior density for a single time slice is

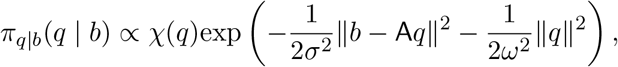

therefore the MAP estimate q_MAP_ is the solution of

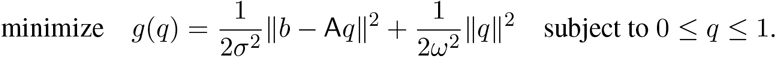

Since this is a quadratic minimization problem with bound constraints, the solution can be found using a projected Newton algorithm [25]. For completeness, we present an outline of how the algorithm proceeds.

We begin by computing the global unconstrained minimizer q^*\**^ of g. Since

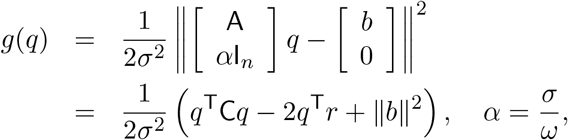

Where

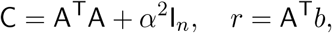

it follows from the first order optimality condition that

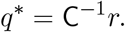

In general, there is no guarantee that the q^*\**^ satisfies the bound constraints, therefore we use it as a starting point for an iterative process leading to a feasible solution. Let

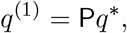

where P is the orthogonal projector onto the unit hypercube, so that

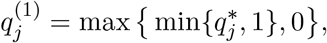

and we identify the set of indices I_act_ *⊂ {*1, …, n*}* corresponding to the components on which the constraints are active, that is, 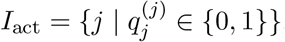, and denote by I_inact_ the set of indices of the components unaffected by the constraints. If I_act_ = *∅*, the global minimizer is inside the feasible set and the algorithm terminates, otherwise we continue as follows. To find the next iterate, we partition the vector q, fixing the components in the active set to the values determined by the projection onto the feasible set. After possibly rearranging the entries of q we have

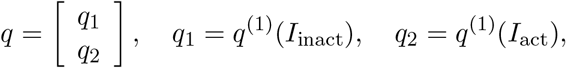

where the components of q_2_ are known and fixed to the extremal values {0, 1}. Next we partition the matrix C and the vector r according to the active/inactive partitioning of q, obtaining

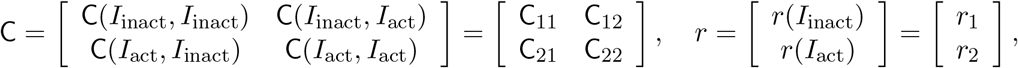

and we write the objective function as

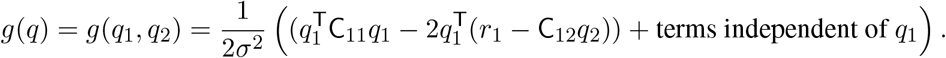

We remark that since q_2_ is constant, the minimizer 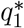 of g(q_1_, q_2_) with respect to the free variables q_1_ is given by

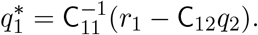

As in the previous step, we project 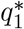 onto the feasible set to guarantee that the constraints are satisfied, obtaining thus the next iterate,

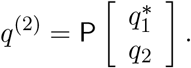

The process continues until no new active components are found. The process is guaranteed to converge in at most *n* steps, although usually it terminates much sooner.

The projected Newton algorithm can be used also to find the MAP estimate for the minimum yank (MY) prior and the mixed prior (MX). In the case of the MY prior, we can write the posterior density as

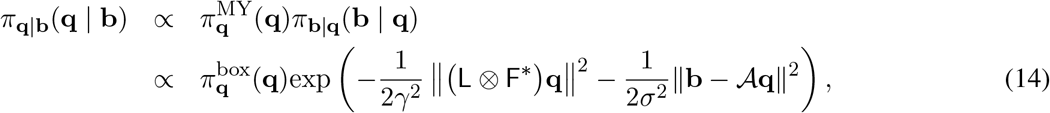

where *A* is the block diagonal matrix (10). Recall that in this case the MAP estimate is a vector **q**^*\**^ = vec(Q^*\**^), with each row of Q^*\**^ *∈* ℒ ^*n×*(*N*+1)^ defining a paths for each muscle, minimizing

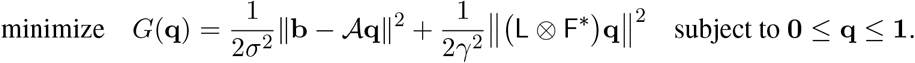

The minimizer can be found with the same algorithm used to find t he M AP w ith t he M A p rior. T he f act that the matrix L *⊗* F^*\**^ has a non-trivial null space may cause technical difficulties. We overcome this shortcoming by assigning boundary conditions at t = t_0_ and/or at t = t_*N*_. Possible choices of boundary conditions will be considered in the next section, where the MCMC sampling from the posterior density is discussed. The same strategy can be adapted for the case of the mixed prior.

### 5.2 MCMC sampling: Independent time slices

We begin by discussing the Gibbs sampler algorithm for the posterior density corresponding to the minimum action prior with the box constraint condition. Because of the mutual independence of the time slices, we limit the discussion to a single time slice, omitting the superscript indicating the time instance. The sampling algorithm was inspired by the engine behind the software Metabolica, designed for analyzing steady state reaction and transport fluxes in complex metabolic networks [11, 12], and later employed to investigate the muscle recruitment problem [27, 28].

After computing the singular value decomposition (SVD) of the lever arm matrix A,

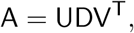

and substituting it in the term describing the equilibrium condition, we obtain

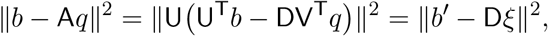

Where

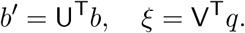

Let r ≤n be the rank of the matrix A, and let d_*j*_, 1 ≤j ≤r be its positive singular values. It follows from the orthogonality of the matrix U that

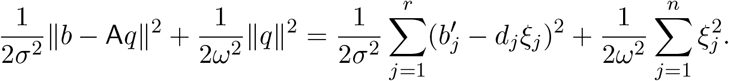

We draw from the posterior using the Gibbs sampler for updating ξ one component at a time. To update the jth component we proceed as follows. Denote by ξ_*c*_ the current value of ξ, and write

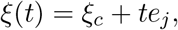

where e_*j*_ is the jth canonical Cartesian unit vector. The random variable ξ(t) is distributed according to a onedimensional Gaussian density with bound constraints. From q(t) = Vξ(t) it follows that

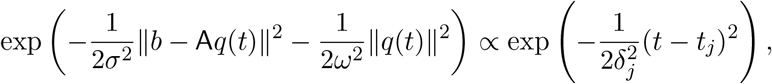

Where

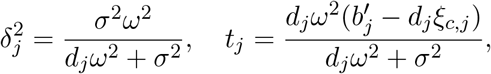

with the understanding that d_*j*_ = 0 if j > r.

To write the bound constraints confining q (t) inside the unit hypercube in terms of t, we observe that t must be chosen so that the inequality

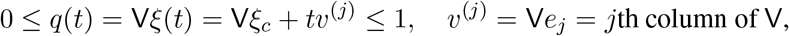

holds component-wise, or equivalently,

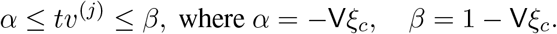

Since t = 0 corresponds to the current value of ξ which satisfies the bounds, the set of feasible values of t is nonempty. To avoid numerical instabilities, we set the components of *v*^(*j*)^ with an absolute value below a threshold τ > 0 to zero, and write the bounds for the components of t as

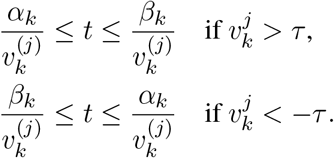

Denoting the index sets where the above conditions hold by J_+_ and J_*−*_, respectively, depending on the sign of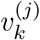, we conclude that

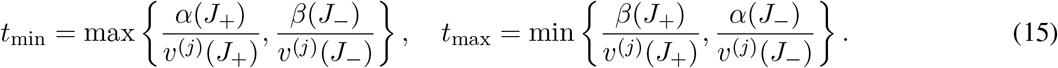

Observe that if the Gaussian is centered at t_*j*_ > t_max_ or t_*j*_ < t_min_, the values of t must be drawn from the tail of the Gaussian. Unfortunately, most of the time the feasible interval is very far in the tail, and the standard error functions cannot be employed. In those cases one needs to resort to numerical quadratures to evaluate the cumulative distribution functions.

Formulas (15) need to be modified when using only the box prior. Formally, the corresponding equations are obtained by the limiting process of letting ω^2^ *→ ∞*. The bound constraints (15) do not change, but the expressions for the mean and variance simplify, giving

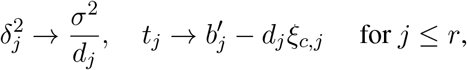

while for r + 1*≤* j *≤*n, the parameter t is drawn from the uniform distribution over the interval [t_min_, t_max_]. We remark that the algorithm for sampling from the posterior with the box prior case is the same as the sampling scheme used in the articles [27, 28].

### 5.3 Sampling of paths and Feynman-Kac model

We turn now to sampling from the posterior densities corresponding to minimum yank and mixed priors, respectively. Unlike in the previous cases, the time slices are no longer mutually independent, thus the longitudinal nature of the priors need to be taken into account.

We consider first the posterior associated with the minimum yank prior (14). Monte Carlo sampling generates an ensemble of paths,

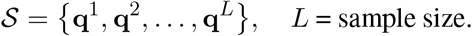

There is a formal similarity of this procedure with the Feynman-Kac path integral formalism of representing solutions of certain diffusion-type parabolic equations as expectations of integrals over paths [13]. In our case the expectation over paths yields the posterior mean estimate of the muscle activation problem with the yank prior. This connection between Monte Carlo sampling and the Feynman-Kac model has been pointed out, e.g., in [7]. It is because of this formal similarity that we refer to the path sampling as Feynman-Kac model.

#### 5.3.1 Fixed boundary conditions

We are now ready to address the question how to generate samples from the posterior density (14). The modifications needed to make it suitable for the mixed prior are outlined at the end of this subsection. Consider first a case in which the boundary values at t = t_0_ and t = t_*N*_ are fixed, 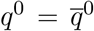 and 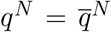. In this case, the time slices q^1^, q^2^, …, q^*N−*1^ are drawn from the density

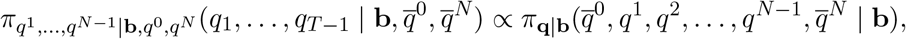

that is, we consider the posterior density with q^0^ and q^*N*^ fixed to the given boundary values.

To generate the sample, we consider a *block Gibbs sampler*, where the updating is done time slice by time slice: Given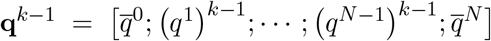, we generate 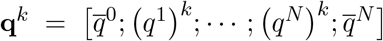 through the following process:

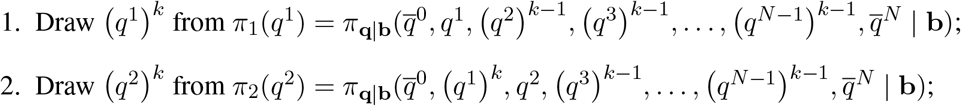

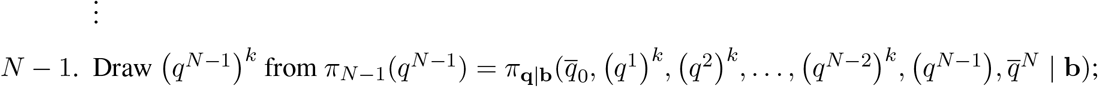

Formally, the task is similar to generating sample paths for the diffusion model corresponding to a *Brownian bridge* [5], a Brownian motion in which the initial and final values are pinned down. The path generation is complicated by the need for the forces to satisfy the the box constraints and the equilibrium conditions. Before discussing these details, consider the term corresponding to the yank term in the posterior density for a single activation vector q^*ℓ*^, 0 < ℓ < N, assuming that all other activation vectors are given. We have

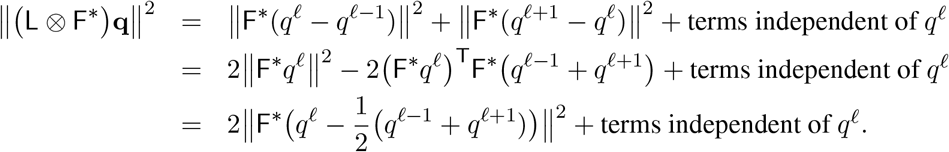

Therefore, ignoring the bound constraints and the equilibrium conditions, the updating of q^*ℓ*^ conditional on the rest of the activation vectors corresponds to drawing from a second order smoothness prior weighted by the matrix F^*\**^ [4], the extra conditions making the process slightly more complicated.

The observations above imply that the ℓth update requires drawing from the density

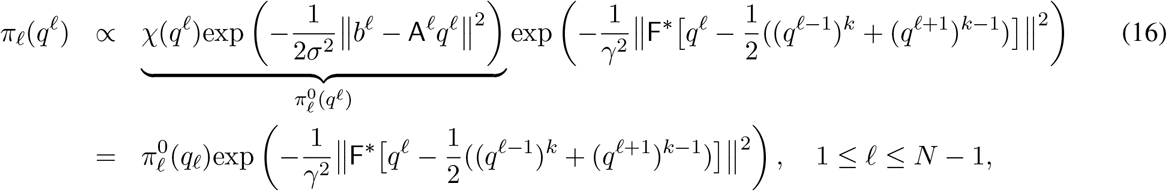

with 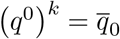 and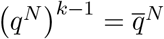.

The block Gibbs sampling algorithm outlined above requires thus the updating of each q^*ℓ*^, keeping the other activation vectors fixed. This can be done as for the updating of the independent time slices, giving rise to the following nested *Gibbs-within-Gibbs algorithm*.

Consider the density (16). Denoting

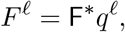

and setting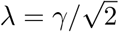, it follows that

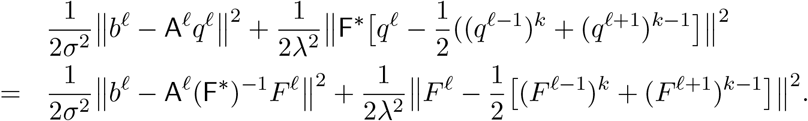

Because the update is computed for one individual time slice at a time, we proceed as in the updating process of independent time slices, using the singular value decomposition of the corresponding equilibrium matrix, scaled by the tetanic forces. Letting

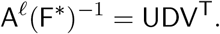

be the SVD of A^*ℓ*^(F^*\**^)^*−*1^, we write the quadratic expression as

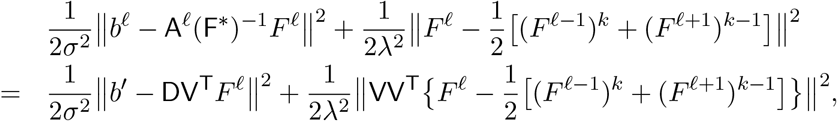

where b^*′*^ = U^T^b^*ℓ*^. Further, letting ξ = V^T^F ^*ℓ*^, we arrive at the expression

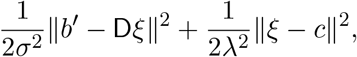

Where

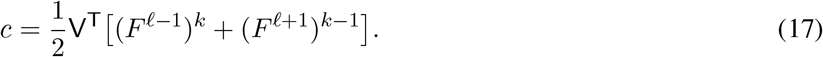

In this manner we have reduced the problem of updating q^*ℓ*^ to the problem of drawing the vector ξ from the distribution proportional to

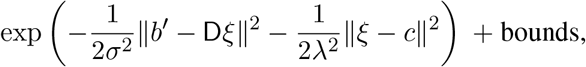

and the process continues component-wise along the lines of the previous subsection.

Sampling the path for the mixed prior is very similar, requiring only the few modifications indicated below. Starting from the formula corresponding to (16),

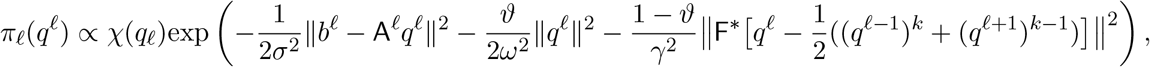

we combine the first two terms in the exponential to obtain

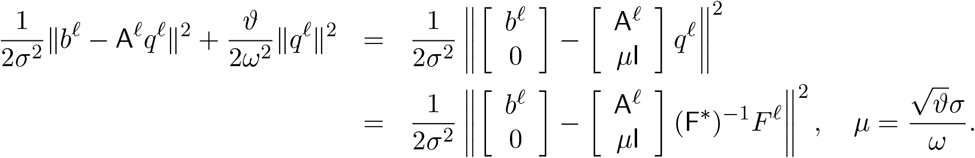

As previously, we compute the SVD,

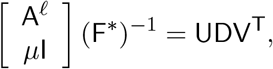

and define

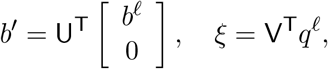

so that the sampling algorithm becomes formally identical to the one presented before.

Above, it was assumed that the initial and final boundary values were given and fixed. Technically, we could fix only one of the two boundary values and treat the other value as a random variable. In general, assuming one or both boundary values known may have a biasing effect on the paths samples. In the rest of this section, we consider some computationally feasible ways to deal with the restriction. In particular, two physiologically meaningful alternatives are proposed.

#### 5.3.2 Periodic boundary conditions

*Periodic boundary conditions* are a meaningful alternative when the dynamics represents a repetitive task, such as level walking: In the last time frame after a completed cycle of steps, the musculoskeletal system is approximately in the same equilibrium position as in the beginning of the cycle, and we may identify q^0^ and q^*N*^ as random variables. Therefore, after initializing the chain by using an initial state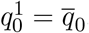, we use the block Gibbs sampler, using in (16) for ℓ = N and ℓ = 0 formally

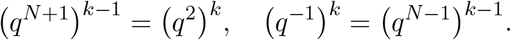

This sampler is similar to a *Brownian ring* of periodic random walks. To avoid ambiguities, the identification of the distributions of q^0^ and q^*N*^ require that in the initial and final equilibrium conditions, the replacements

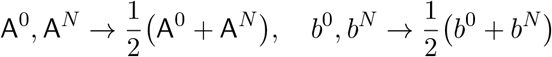

are performed before the sampling.

#### 5.3.3 Freeing the fixed boundary values

Another possibility to free the fixed boundary conditions is to postulate prior distributions for the end values, and write

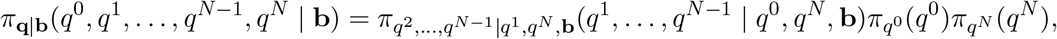

where 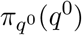 and 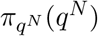 represent the initial and final distributions.

In practice, we draw the new independent initial and final values from their respective distributions before each block Gibbs updating round. While this extension removes the necessity to specify exactly the boundary values, an initial distribution needs to be entered, which may also bias the outcomes.

### 5.4 Augmenting the likelihood

The main focus of the discussion above was on prior models, while the likelihood was restricted to contain information about the equilibrium condition only. Sometimes, however, we may have additional observations available, such as EMG-recordings of a given set of muscles. While the EMG data do not translate directly to muscle activation (see [2] for a discussion), they carry indirect information that should be taken into consideration in the analysis [34]. Consider, for simplicity, EMG data collected from one single muscle. The most straightforward model is to assume a strictly increasing dependency between the rectified and low-pass filtered EMG recording μ(t) of the jth muscle and the activation level of the muscle,

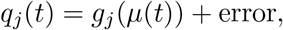

where g_*j*_ is an appropriately chosen increasing function like a sigmoid, and the “error” refers to a modeling error. A computationally feasible approach is to use the activation model above to define a likelihood interval, requiring that q_*j*_ (t) satisfies

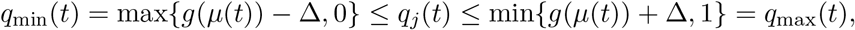

where Δ > 0 is the presumed model accuracy. This model translates into the likelihood model by multiplying the likelihood density (11) by the indicator functions of the intervals [q_min_(t_*ℓ*_), q_max_(t_*ℓ*_)],

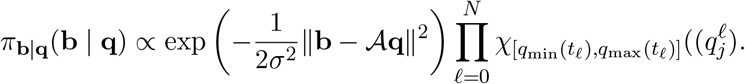

This modification can be effectuated by simply replacing the lower and upper bounds of 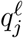 in the box prior (12) by the new bounds, thus the modification of the Gibbs sampler algorithm becomes straightforward.

## 6 Computed examples

In this section, we demonstrate the effect of different priors and selections of boundary conditions and parameters.

The generic musculoskeletal model Gait2392 [9] was employed in this study to simulate level ground walking. The model, illustrated in Figure 1, was scaled to closely approximate the anthropometry of a 86 year old man with an instrumented knee prosthesis, and includes 2 lower limbs and the torso, comprising of total of 12 bodies from the torso down to the toes, 92 actuators representing the muscles, 46 for each leg, and 17 degrees of freedom: 3 at the hip, one at the knee, ankle, subtalar joint, and metatarsophalangeal joint for each leg, and 3 between the pelvis and the torso. The experimental data are part of the fifth dataset of the Knee Grand Challenge [10]. The biomechanical variables were estimated using OpenSim, an open-source platform to perform biomechanical analyses of musculoskeletal dynamics [8]. Joint angles and joint torques were computed by implementing and solving the inverse problem, as described in Section 2. The muscle moment arms defining the moment matrices A^*ℓ*^ were obtained through the Analyze Tool of OpenSim once the joint angles were known.

**Figure 1:**
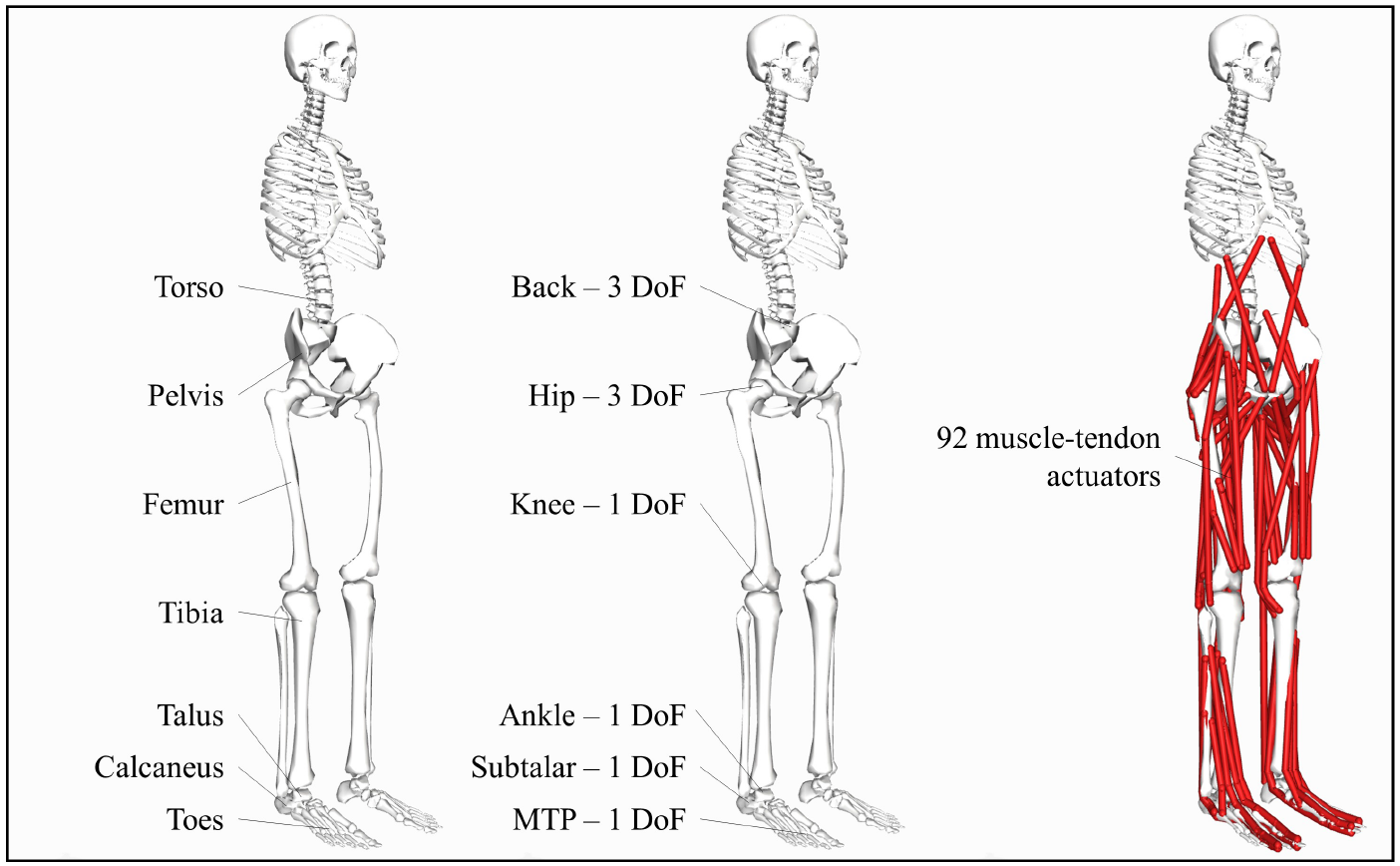
OpenSim model employed in current simulations. The model consists of 12 bodies divided into two lower limbs and a torso as indicated in the figure, 17 degrees of freedom and 92 muscle-tendon actuators, 46 for each leg.

As a starting point, we assume that the inverse dynamic problem is thus completely solved, and the lever arm matrices and the corresponding torque vectors 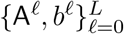, where A^*ℓ*^ *∈* ℒ ^*m×n*^, b^*ℓ*^ *∈* ℒ ^*m*^, are resolved with accuracy that allows us to use an equilibrium model (4) with noise modeled as

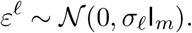

In our data set, the number of equilibrium equations is m = 17, while the number of muscles included in the model is n = 92. The time interval of the full stride is divided in L = 130 intervals. Observe that the vectors b^*ℓ*^ represent generalized forces, and in order to scale the standard deviation properly, we write

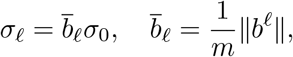

and set σ_0_ = 0.001.

We start the simulations by running the MCMC sampler using the box prior, i.e., including only the physiological bound constraints to muscle activations, and subsequently we compute samplers for three more informative priors, minimum activation (MA), minimum yank (MY) and mixed (MX) priors. In all simulations, the sample size is set at N = 100 000, while the prior and likelihood variance parameters are set as indicated in Table 1. The results are summarized in figures that show quantile plots as described in detail below. The plots are limited to only eight muscles, four large ones (*Gluteus Medius Anterioris L, Medial Gastrocnemius L, Soleus L*, and *Tibial Anterioris L*) and four smaller muscles (*Sartorius L, Gracilis L, Flexor Digitorum L*, and *Extensor Digitorum L*). The locations of the muscles are shown in Figure 2

**Table 1:** The values of the parameters used in the simulations. In all simulations, the sample size is N = 100 000.

| Prior | $\sigma_0$ | $\omega$ | $\gamma_0$ | $\vartheta$ |
| --- | --- | --- | --- | --- |
| Box | 0.001 |  |  |  |
| Min. activation | 0.001 | 0.1 |  |  |
| Min. yank | 0.001 |  | 0.01 |  |
| Mixed | 0.001 | 0.1 | 0.01 | 0.5 |

**Figure 2:**
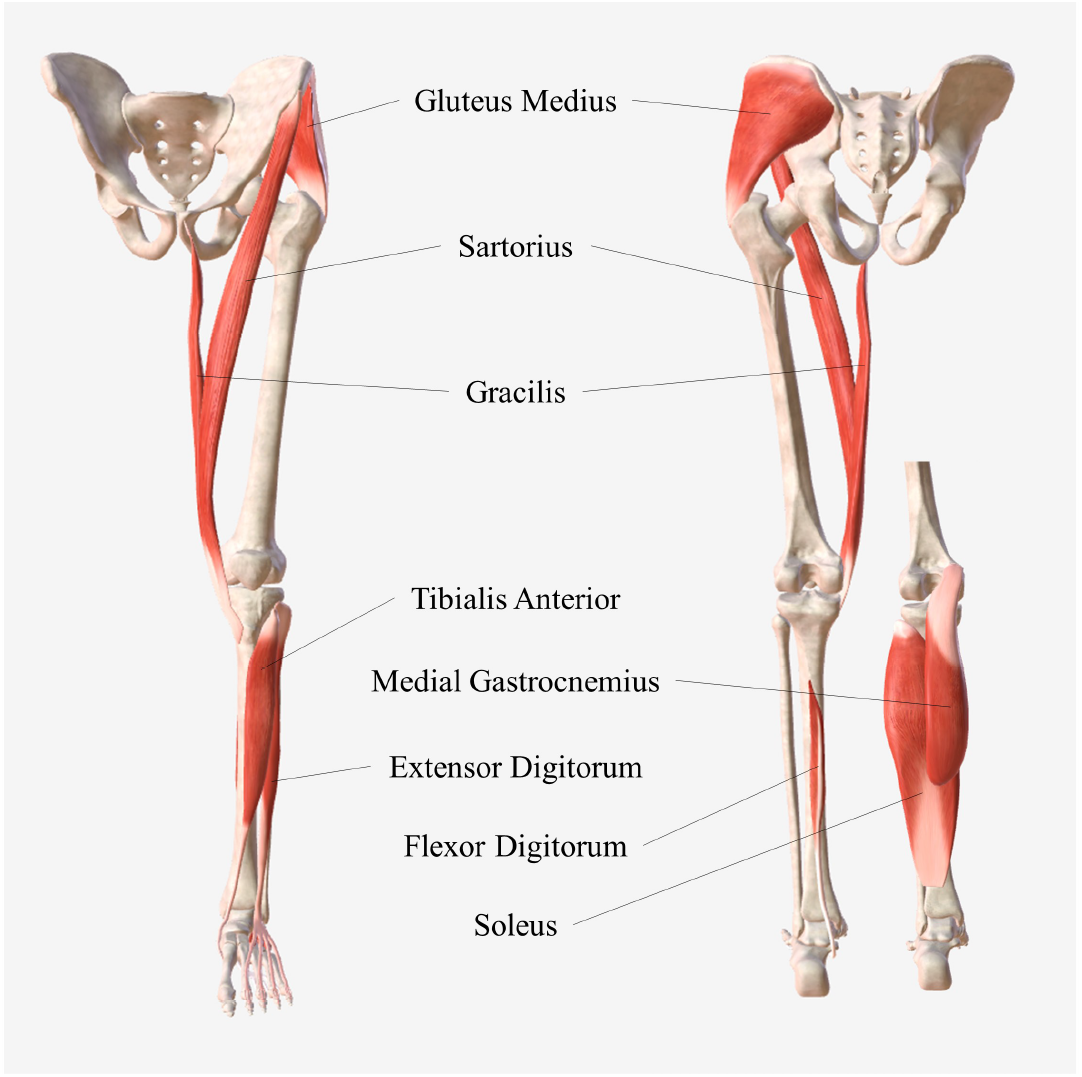
The eight selected muscles of the left leg whose activation patterns are shown in this study. Anterior (left) and posterior (right) view.

Figure 3 shows a superposition of the results corresponding to the MA box prior. In the figure, the interval between the minimum and maximum sample value at each time slice of the box prior is indicated by a thin red bar, while the interval of the 75% quantile is plotted with a thick red bar. On top of that, we have plotted the envelopes containing 75% (darker shade) and 100% (lighter shade) of the sample points corresponding to the MA prior sampling. Finally, the median of the MA sample is indicated by a blue curve, and the MAP estimate computed by the optimization algorithm is plotted in red. Using the same color coding, the results corresponding to the smaller muscles are shown in Figure 4.

**Figure 3:**
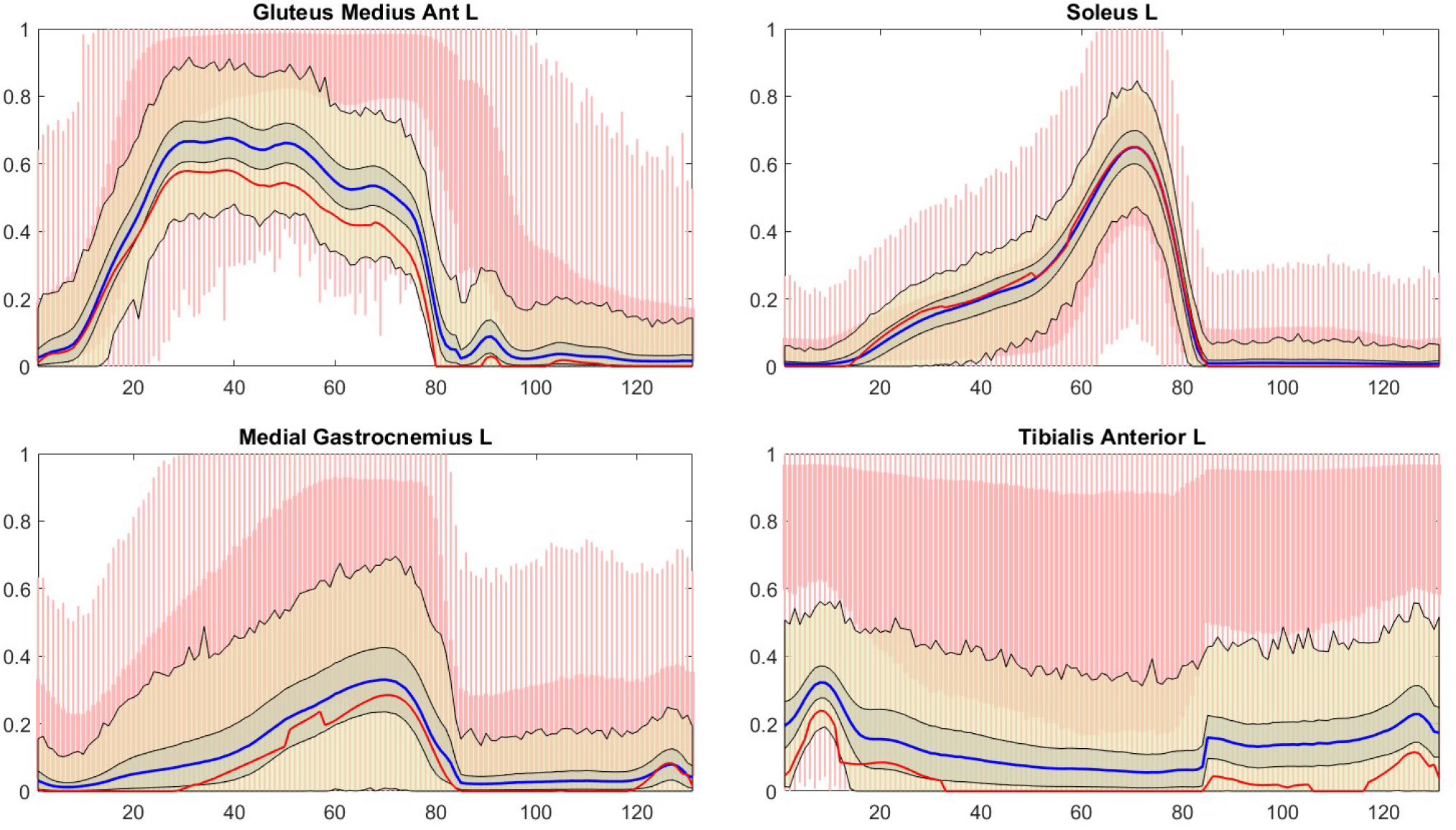
Sampling results corresponding to the MA prior. The thick red bars correspond to the 75% quantile of samples computed with the box prior, while the thin red bars indicate the full range of the samples. Similarly, the dark and light beige envelopes correspond to the 75% quantile and the full range, respectively, of the sample based on the MA prior. The red curve is the computed minimum activation MAP estimate, and the blue curve is the median over the MA sample. The sampling time step in all figures is Δt = 0.00833 s, the total time interval being therefore about 1.1 seconds.

**Figure 4:**
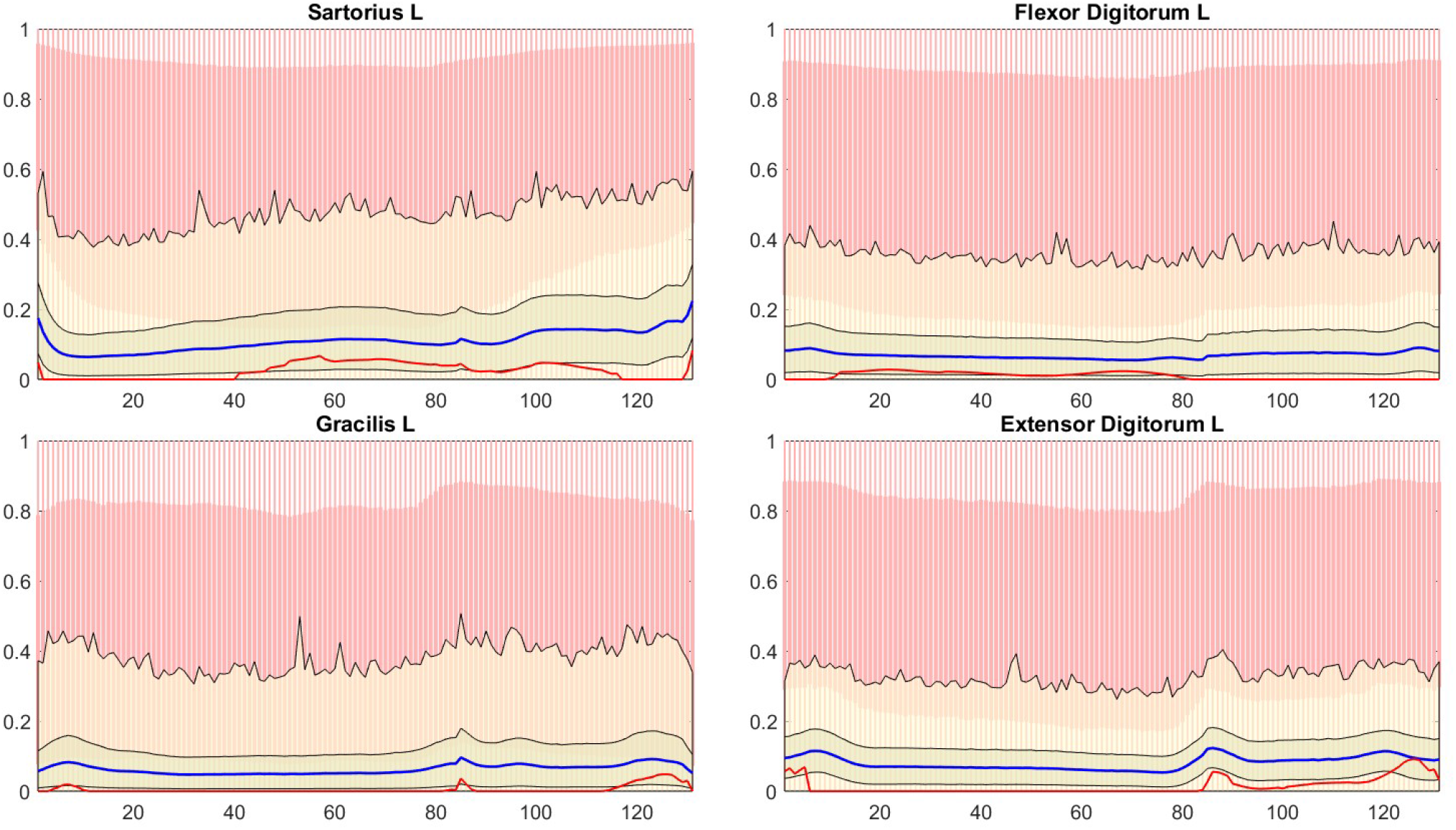
The sampling results corresponding to Figure 3 for the selected smaller muscles. The color coding is as in the previous figure.

The results indicate that as expected, the MA prior drives the sample envelope, and the MAP estimate in particular, to the low end of the box prior feasible intervals. In particular, the samples of the smaller muscles seem to reflect very weakly the performed task of level walking, showing an almost flat response.

We then run the sampler using the minimum yank prior. The standard deviation γ is given in the units of forces (N), the implicit dependency of the length of the time step being suppressed, and in order to have dimensionless parameters, we scale γ by the average of the tetanic forces, writing.

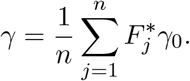

To set the value γ_0_, literature about the rate of force development (RFD, [N/s]) or rate of torque development (RTD, [Nm/s]) was consulted [15, 23, 32, 35]. The RFD and RTD parameters represent the ability of a muscle to rapidly develop external force. Based on the experimentally measured values, we defined a plausible range for the yank. Subsequently, to obtain a plausible value for γ, the value range of the yank, originally estimated for one second, was scaled to the time step of the example data, corresponding to Δt = 0.00833 s. The value of γ_0_ used in the simulations is given in Table 1.

In the first simulation, the initial and final activations patterns, q^0^ and q^*N*^, are fixed to be equal to the median values of the corresponding activations of the box prior sample. Figures 5 and 6 show the envelopes containing 75% of the sampled values as well as the full intervals of the values. For comparison, the envelopes are superimposed on the intervals containing the full intervals of the sampled values using the box prior and the minimum activation priors, respectively, giving an idea of how much the minimum yank prior restricts and translates the values.

**Figure 5:**
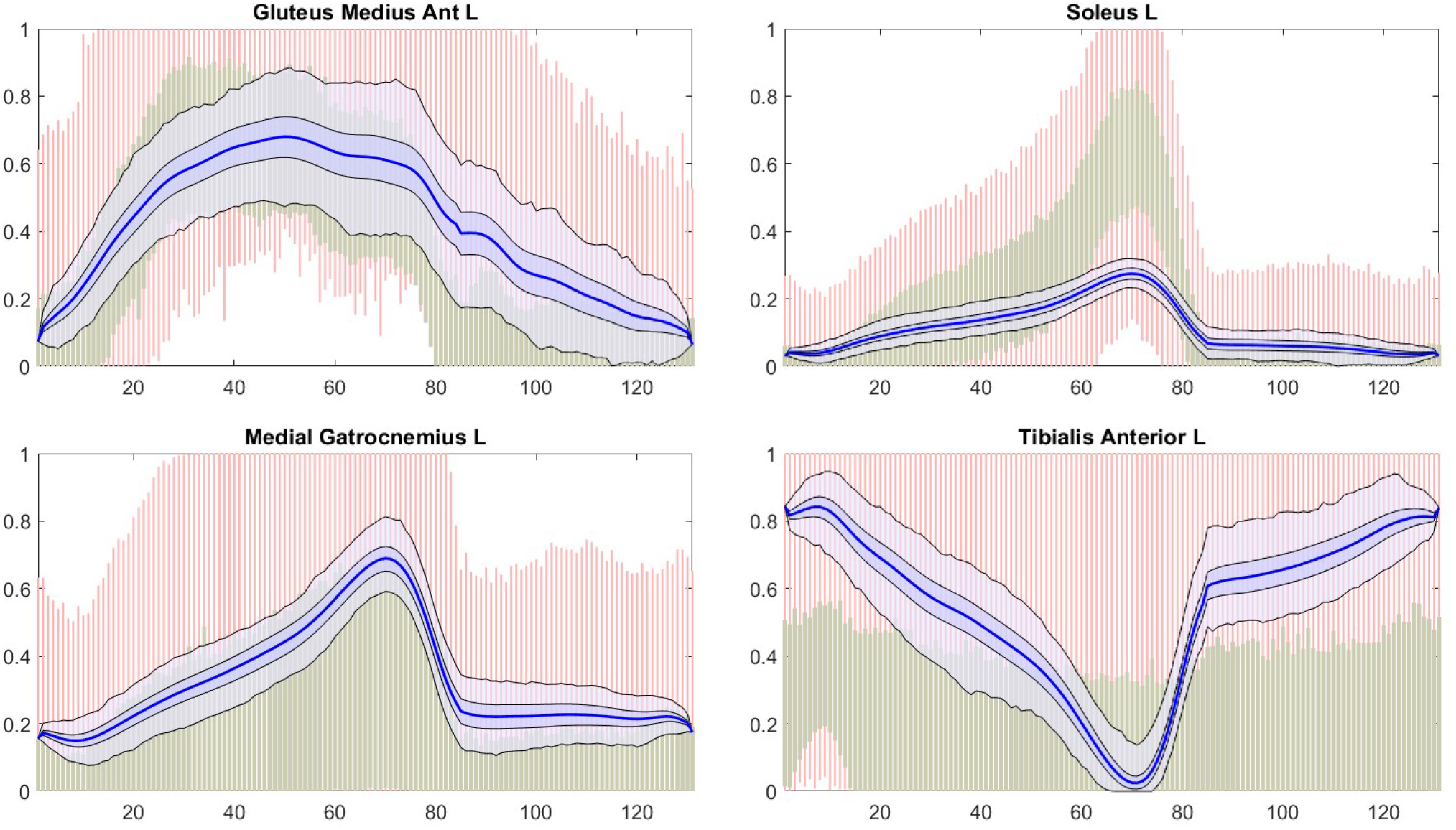
Sampling results corresponding to the minimum yank prior with fixed boundary conditions, the boundary values being fixed at the median value of the sample drawn with the box prior. The red bars indicate the full range of the sample using the box prior, and the green bars correspond to the full range corresponding to the minimum activation prior. The dark and light blue envelopes correspond to the 75% quantile and the full range of the paths drawn with the MY prior, respectively. The blue curve is the median over the MY sample.

**Figure 6:**
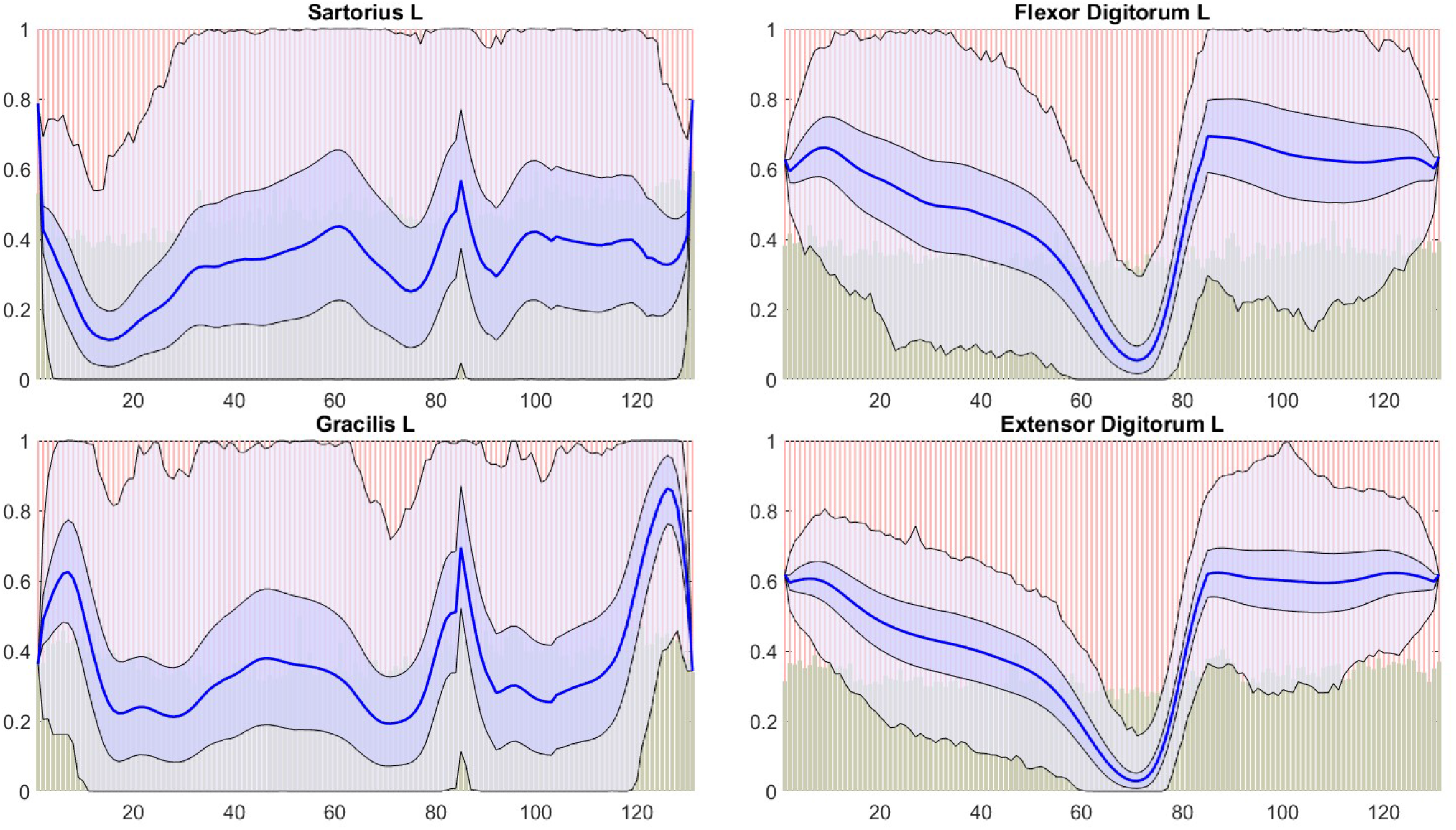
The figure corresponding to Figure 5 for the selected smaller muscles. The color coding is as in the previous figure.

To investigate the effect of the fixed boundary conditions on the envelopes, for comparison we run the sampler using the periodic boundary conditions. The results are shown in Figure 7 for the selected larger muscles and in Figure 8 for the smaller ones. The results with the smaller muscles, in particular, indicate that the periodic boundary condition may be overly lax, leaving a large uncertainty in the muscle paths.

**Figure 7:**
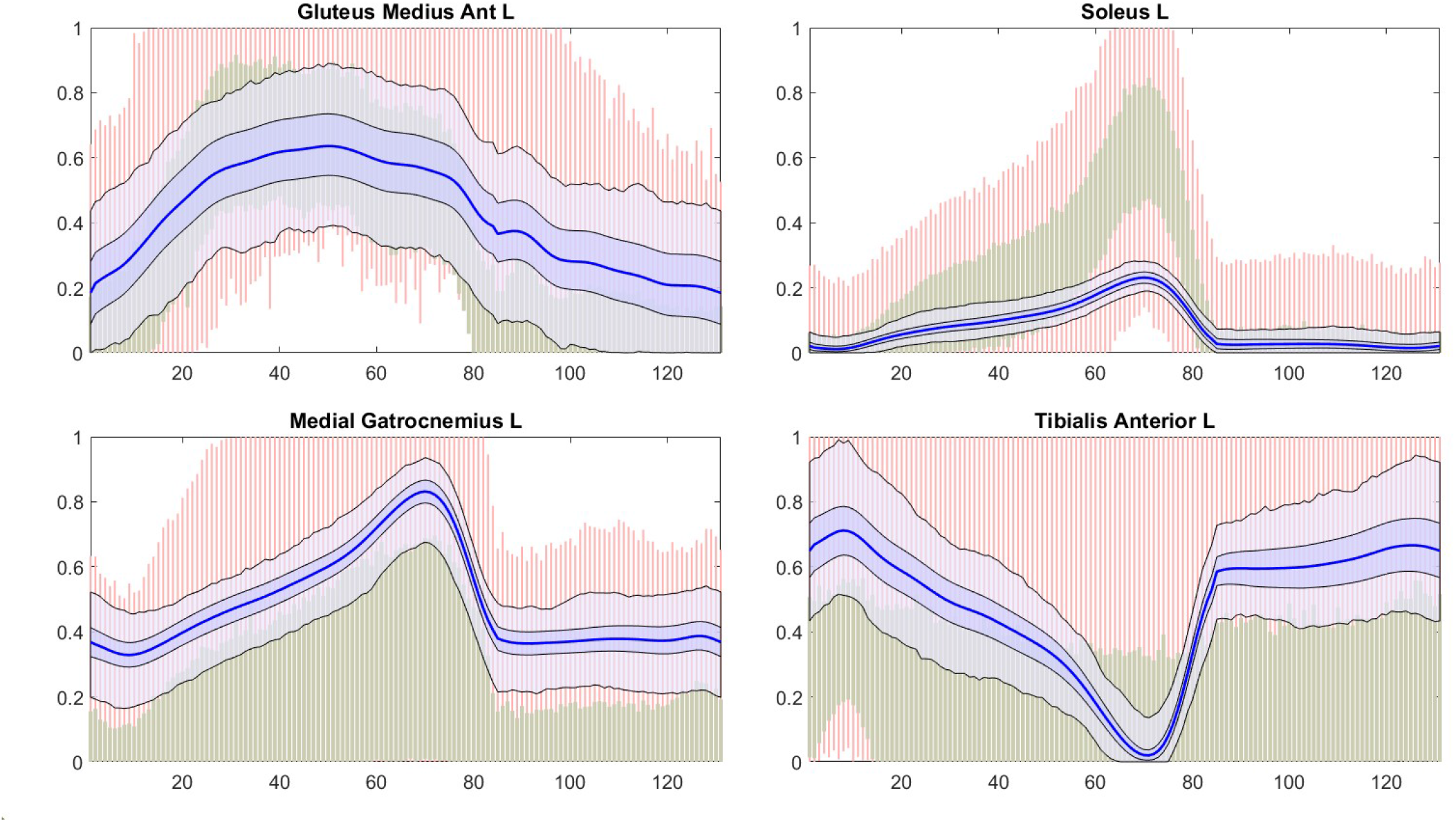
Sampling results corresponding to the minimum yank prior with periodic boundary conditions. The red bars indicate the full range of the sample using the box prior, and the green bars correspond to the full range corresponding to the minimum activation prior. The dark and light blue envelopes correspond to the 75% quantile and the full range of the paths drawn with the MY prior, respectively. The blue curve is the median over the MY sample.

**Figure 8:**
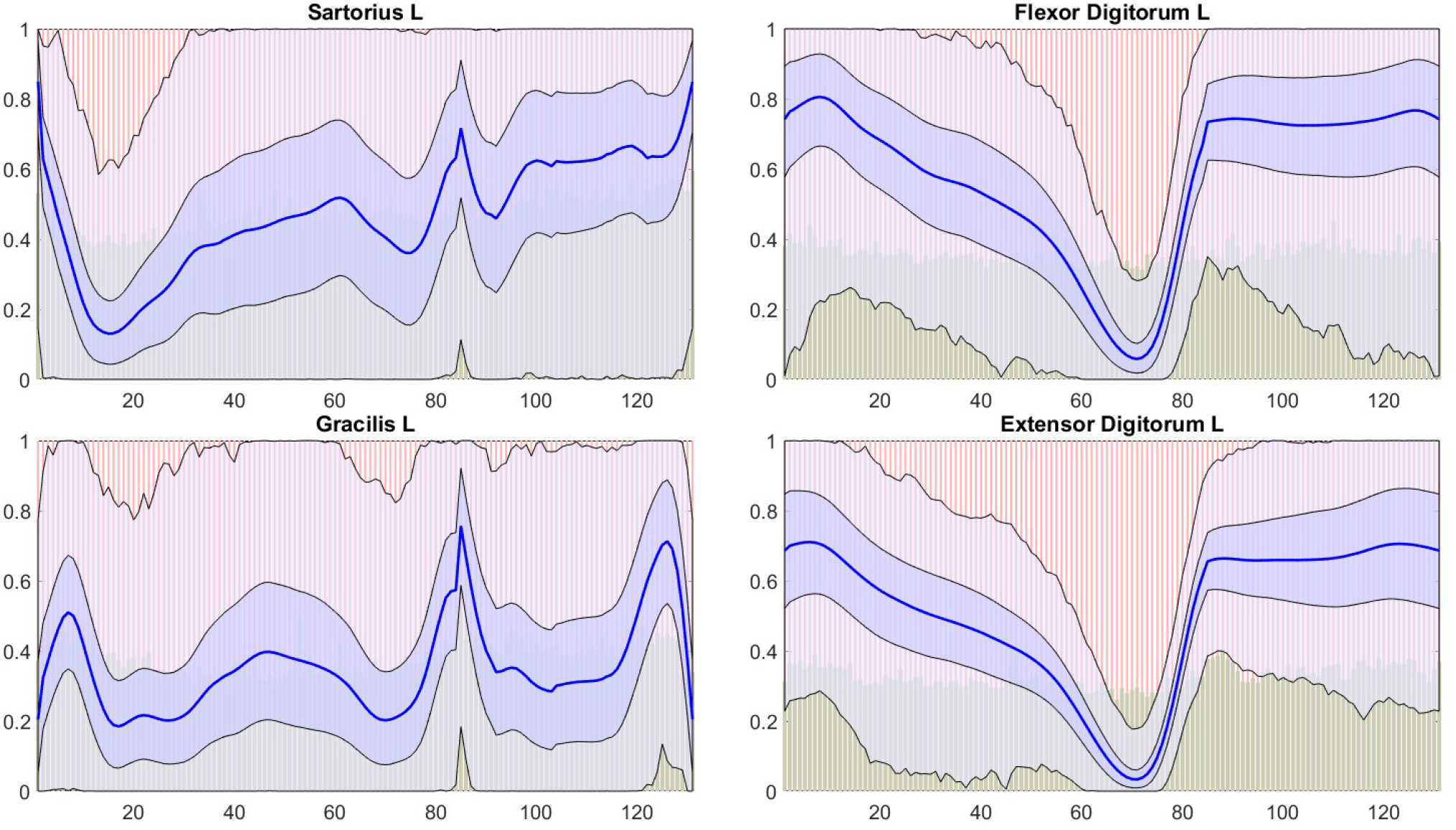
The figure corresponding to Figure 7 for the selected smaller muscles. The color coding is as in the previous figure.

Finally, we generate the samples using the mixed prior, setting the value of the interpolation parameter to ϑ = 1/2. Figures 9 and 10 show the envelopes corresponding to the fixed boundary conditions. Here, the initial and final values were set to be equal to the median values of q^0^ and q^*N*^ over the sample corresponding to the minimum activation prior. Again, the envelopes of 75% quantile and full range are plotted against the full range of the box prior (red bars) and the minimum activation prior (green bars). For comparison, the corresponding plots using the periodic boundary condition are shown in Figures 11 and 12, respectively.

**Figure 9:**
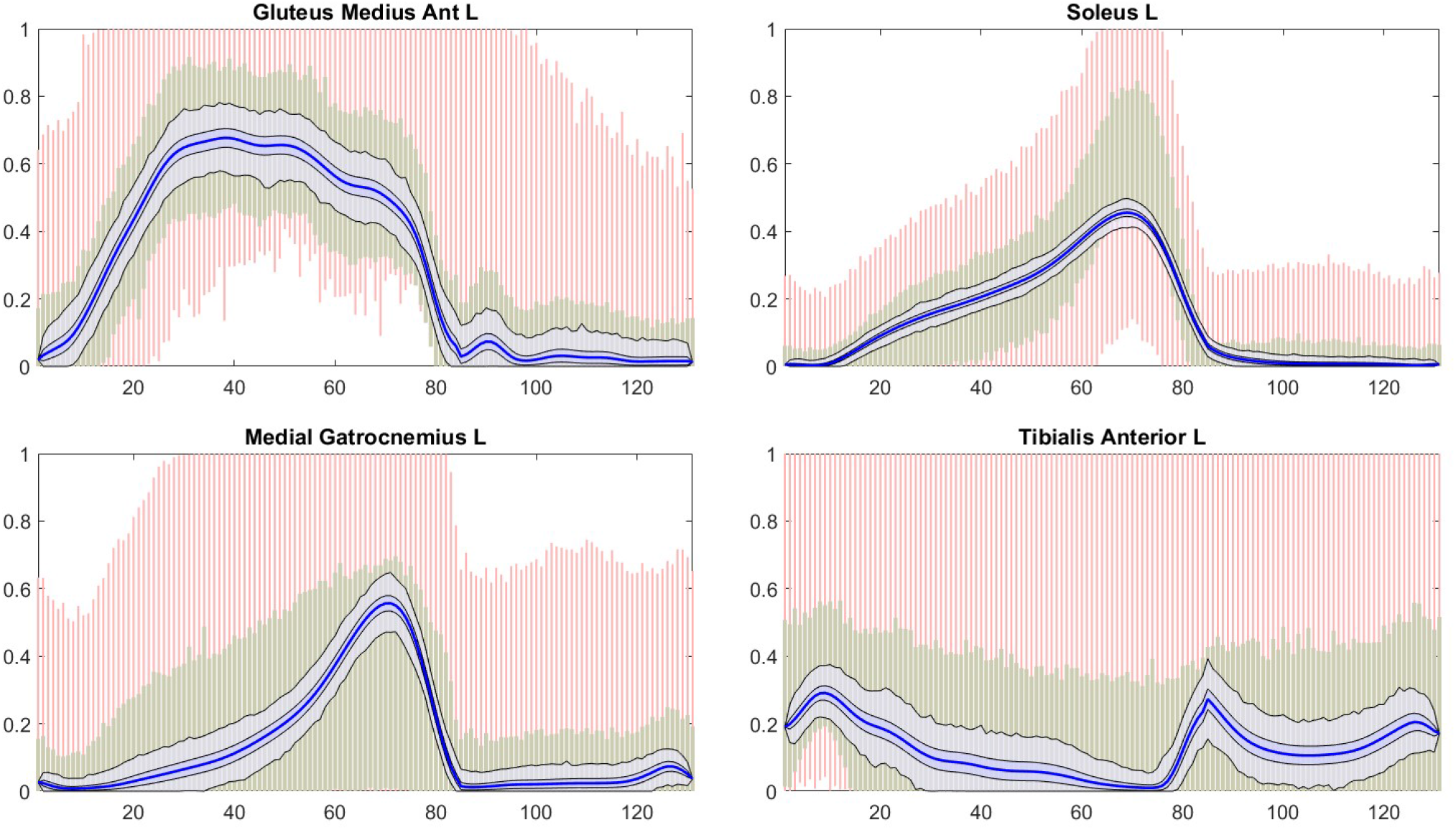
Sampling results corresponding to the mixed prior with fixed boundary conditions, the boundary value as before fixed at the median value over the minimum activation prior sample. The red bars indicate the full range of the sample using the box prior, and the green bars correspond to the full range corresponding to the minimum activation prior. The dark and light blue envelopes correspond to the 75% quantile and the full range of the paths drawn with the MX prior, respectively. The blue curve is the median over the MX sample.

**Figure 10:**
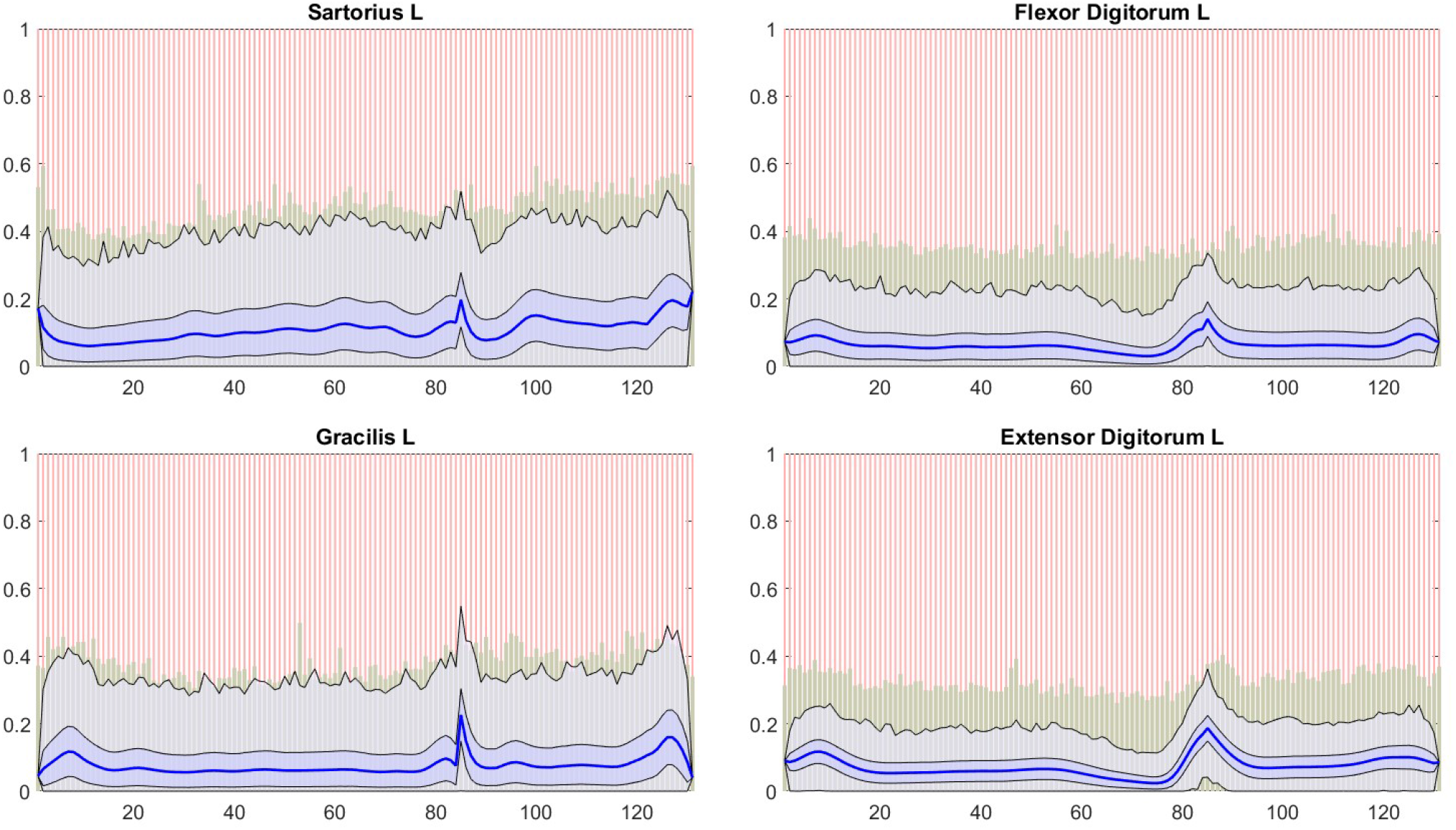
The figure corresponding to Figure 9 for the selected smaller muscles. The color coding is as in the previous figure.

**Figure 11:**
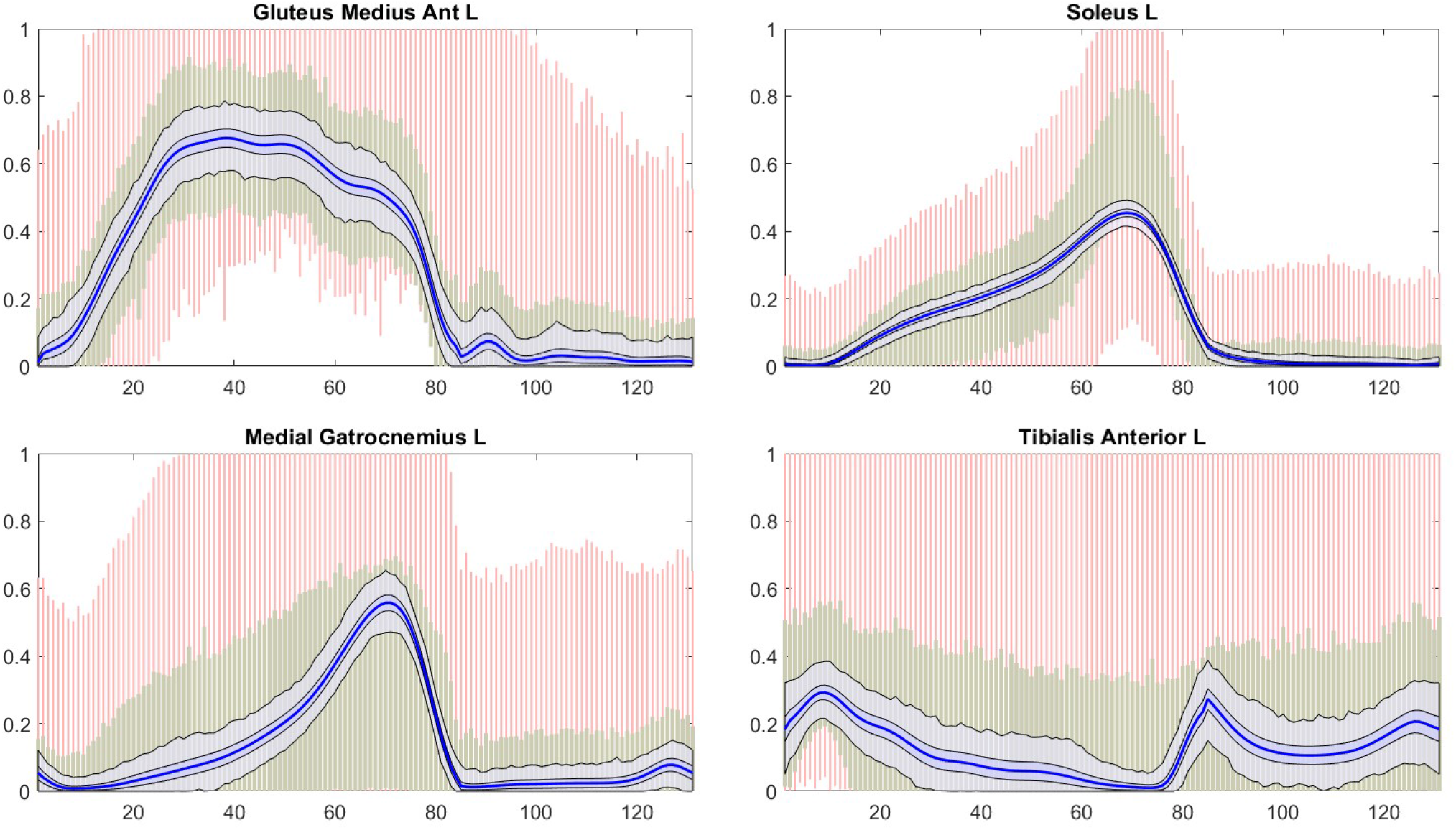
Sampling results corresponding to the mixed prior with periodic boundary conditions. The red bars indicate the full range of the sample using the box prior, and the green bars correspond to the full range corresponding to the minimum activation prior. The dark and light blue envelopes correspond to the 75% quantile and the full range of the paths drawn with the MX prior, respectively. The blue curve is the median over the MX sample.

**Figure 12:**
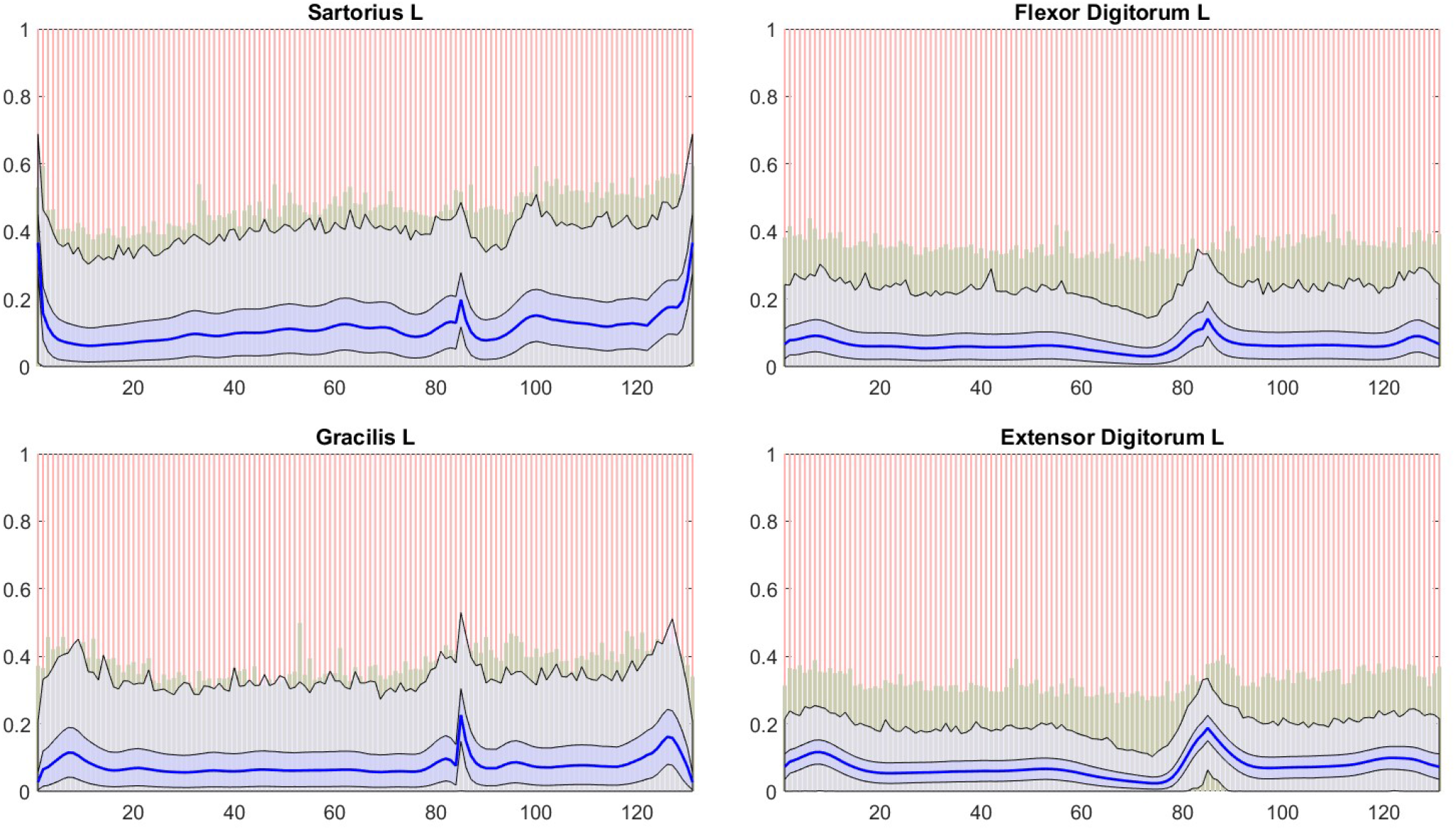
The figure corresponding to Figure 11 for the selected smaller muscles. The color coding is as in the previous figure.

## 7 Discussion

The Bayesian framework provides a rich environment of investigating the muscle recruitment problem by sampling the probability distributions representing possible activation configurations that respect the equilibrium conditions up to a prescribed accuracy. The previous works [27, 28] were limited to consider equilibrium solutions with the minimum prior assumption that the natural bound constraints were satisfied with no other preferential properties given. Here, the formalism is extended to include further prior information. Unlike the minimum activation prior that still assumes the different time slices to be independent, the minimum yank and mixed priors introduce longitudinal prior structure, tying together the time slice realizations through a smoothness prior of first order. While the minimum activation prior can be justified, e.g., through energetic considerations, the longitudinal prior structure is meaningful as the musculoskeletal system under normal conditions and outside the regime of extreme movements is likely to favor smooth paths without jerky activations that may even be harmful for the muscles. Considering the results, significant differences in the activation patterns can be seen both in the large and small muscles. For instance, comparing the activation pattern of the soleus muscle in Figure 3 on one hand and in Figures 5 and 7, we see that the minimum activation prior allows a significantly higher activation level than the minimum yank prior, regardless of the boundary condition, indicating that while the minimum activation prior aims overall to favor lower activation levels, in single muscles, the desired effect is achieved by restricting the yank. Conversely, comparing a smaller muscle such as extensor digitorum muscle in Figures 4 with those in Figures 6 and 8, the minimum yank prior leads to a higher activation level. In fact, the minimum activation prior seems to favor solutions that are close to zero, which, while possible from the point of view of the equilibrium, may not correspond to a typical activation during normal level walk, see, e.g, the supplementary EMG data in [16], and foot EMG data in [18]. These considerations promote models with longitudinal priors, however, further evidence to definitely justify the use of one prior over another is required. Future tests for model validation include a comparison of the joint contact forces predicted by the models based on the muscle forces to the in vivo recordings from the instrumented implants.

A standard deterministic approach to solve the muscle recruitment problem is to use optimization tools to find a single trajectory that minimizes or maximizes a given objective function. The Bayesian formulation described here can be used to produce such optimal solutions by maximizing the posterior density, or, equivalently, minimizing the corresponding Gibbs energy, defined as the negative logarithm of the posterior probability density. Formally, the Gibbs energy corresponds to a penalized least squares solution with bound constraints. In particular with the minimum yank prior, a caveat concerning the MAP estimate is in order. The MAP estimate is easy to find by using the projected Newton algorithm described in this paper. However, while the minimum yank prior favors smooth solutions with no sudden jerks, the hard boundary conditions may make the MAP estimate to be non-smooth: When the smooth solution meets the boundary, typically an abrupt flattening of the curve ensues, creating a sudden kink that does not respect the smoothness requirement. For this reason, the MAP estimate does not necessarily represent the typical paths. We point out that there is also a physiological reason to assume that the muscles anticipate the change in the activation state: Human walking is sometimes characterized as “controlled falling”, and the muscles secure the joints through co-contractions before the reactive forces activate. The sampled solutions with the MY and MX priors do qualitatively demonstrate this behavior. The fact that the MAP estimate may not be a good representative of the density is nothing new in the Bayesian context, which is the reason why the computations of MAP estimates are sometimes not recommended in the literature.

## Acknowledgement

The work of MA and ES was partly supported by the NSF grant DMS 2204618, and DC by the NSF grant DMS 1951446.

